# Immobilized dicot and monocot viral vectors enable rapid screening of RNA mobility elements for mobile RNA engineering and RNA-based genome editing

**DOI:** 10.64898/2026.07.31.741600

**Authors:** Nathaniel M. Butler, Charlene M. Grahn, Colby Starker

## Abstract

RNA mobility has emerged as a valuable component of RNA-based genome editing and DNA-free transformation technologies. However, experimental systems for rapidly evaluating RNA mobility remain limited, particularly in monocot species where grafting approaches are not feasible. Here, we developed immobilized versions of *Foxtail Mosaic Virus* (FoMV) and *Tobacco Rattle Virus* (TRV) with impaired systemic viral movement as generalizable platforms for transient expression and functional screening of mobile RNAs. A simple *Nicotiana benthamiana* leaf assay enabled direct visualization and molecular quantification of transcript mobility using fluorescent reporter fusions carrying seven previously described RNA mobility elements from dicot and monocot species. The platform consistently distinguished mobile elements displaying higher or lower frequencies of mobility across both viral systems, with T-RNA-like sequence (TLS), TLSgly and maize *FLOWERING LOCUS T* (*FT*) ortholog, ZCN19, and as well as ZCN16 displaying significantly higher frequencies of mobility compared to non-mobile element controls in FoMV and TRV, respectively. Translation of these findings to virus-induced genome editing demonstrated that mobile elements identified through the screening platform enhanced FoMV-mediated editing of *PHYTOENE DESATURASE* in *Setaria viridis* (*SvPDS*), with TLSgly increasing somatic editing frequencies approximately two-fold relative to sgRNA alone. Together, these results establish immobilized FoMV and TRV platforms as versatile screening tools for evaluating RNA mobility, optimizing RNA cargos for viral genome editing, and a scalable framework for engineering mobile RNAs and accelerating development of RNA-based technologies for functional genomics and crop improvement.

## Introduction

RNA mobility and cell-to-cell movement of mRNA in plants has gained new significance with the development of RNA-based gene editing and morphogenic regulator-mediated transformation systems. Traditional methods of plant transformation have created a so-called “bottleneck” for functional genomics studies and development of improved crop varieties using modern genetic engineering tools, such as CRISPR (Wu et al., 2026). For this reason, alternative methods of plant transformation have been developed utilizing morphogenic regulator genes, such as *isopentenyltransferase* (*ipt*) and *Babyboom* (*Bbm*)-*Wuschel2* (*Wus2*) to create transgenic organs or embryos by induction of de novo meristematic tissues in both whole plants and tissue culture explants (Butler et al., 2025; Lowe et al., 2016; Youngstrom et al., 2024). Although each method provides different benefits, they largely share the common reliance of incorporation of new DNA to be most effective.

DNA or transgene-free gene editing (referred to herein as DNA-free editing) is a genetic engineering approach associated with the delivery of gene editing reagents (i.e. Cas-protein and sgRNA) without the incorporation of new DNA. Reducing reliance on incorporation of transgenes for editing eliminates the need to excise editing reagents or generate null segregates and facilitates editing in species that cannot effectively support these systems (Bhattacharjee et al., 2023; Cai et al., 2025; Su et al., 2026). There are also important regulatory and off-target effect considerations since transgene integration events occur randomly in the genome and are regulated in most counties (Domingo, 2025; Turnbull et al., 2021). Hence, a number of DNA-free gene editing approaches have emerged including delivery of pre-assembled gene editing reagents to protoplasts or explant tissues via particle bombardment or nanoparticles (Svitashev et al., 2016; Y. Zhang et al., 2021, 2022), or RNA-mediated approaches utilizing plant RNA viruses and mobile RNAs (Nagalakshmi et al., 2022; Park et al., 2026; Yang et al., 2023). RNA-mediated approaches largely have a reduced reliance on plant transformation and tissue culture but variable effectiveness for generating heritable edits.

The study of RNA mobility in plants began with investigating mobile mRNA transcripts, such as maize *KNOTTED1* and Arabidopsis *FLOWERING LOCUS T* (*FT*) and associated functions (Li et al., 2009; Lucas et al., 1995). A fusion of the tomato *KNOTTED1*-like (*LeT6*) and *PYROPHOSPHATE-DEPENDENT PHOSPHOFRUCTOKINASE* (*PFP*) transcripts (*PFP-LeT6*) in grafted tomato plants led to drastic alternations in distal leaf morphology (Kim et al., 2001) while mobility of the Arabidopsis *FT* transcript shed light on a vital long-distance signaling system utilized by most vascular plants (Lu et al., 2012). Further investigations into the genetic features underlying mobility of these transcripts and others have led to the discovery of *cis*-acting RNA mobile elements, such as T-RNA-like sequences (TLSmet, TLSmetΔDT, TLSgly, TLSIle; Zhang et al., 2016) and truncated forms of the Arabidopsis FT transcript (AtFTtruc; Lu et al., 2012) and maize orthologs (ZCN16 and -19; Beernink et al., 2022). The majority of these discoveries have been made through transcript fusions and grafting studies. However, these methods are technical and laborious, requiring the creation of transgenic lines and testing using species capable of grafting (Thomas & Frank, 2019). These restrictions have severely limited high-throughput validation of mobile transcripts and the study of RNA mobile elements in monocot species.

Plant RNA viruses, such as *Foxtail Mosaic Virus* (FoMV), have created new opportunities for transient gene expression without the incorporation of new DNA and studying mobile RNA (Beernink & Whitham, 2023; Bouton et al., 2018). FoMV and other members of the Potexvirus genus, including *Potato Virus X* (PVX) have monopartite, positive-strand RNA genomes which are amenable to Agrobacterium-mediated delivery on a T-DNA backbone. Potexviruses are highly infections and can spread systemically through host vascular tissues via encapsulation by coat proteins (CPs) and cell-to-cell movement through plasmodesmata by movement proteins (MPs) and other viral factors (Verchot-Lubicz et al., 2007). PVX and other viruses have been immobilized by mutagenesis of these viral factors, reducing their virulence and cell-to-cell movement (Yu et al., 2020). This approach has provided opportunities to transiently express and study RNAs encoded in the viral genome without systemic mobility of the virus and could be extended to other dicot viruses outside of Potexviruses, such as *Tobacco Rattle Virus* (TRV).

In this study, FoMV and TRV genomes were mutagenized to restrict cell-to-cell movement and optimized for transient expression of a fluorescent marker, AmCyan and sgRNAs to test the function of RNA mobile elements in both dicot and monocot systems. Both immobilized viral systems demonstrated transient expression restricted to infected, source tissues with evidence of RNA mobility occurring with the introduction of mobility elements. Although similar trends were observed across the two viral expression systems, differences were observed – predominately between RNA mobility elements originating from monocots versus dicots. These data were validated in whole-plant infection experiments, demonstrating enhanced somatic viral editing in setaria (*Setaria virdis*). This study provides important resources for testing RNA mobile elements and optimizing RNA reagents for plant gene editing and morphogenic regulator-mediated transformation.

## Results

### Immobilizing FoMV and TRV plant RNA viruses

Previous characterizations of FoMV and TRV genomes have provided open reading frames (ORFs) and annotated protein functions as candidates for immobilization via mutagenesis (Beernink & Whitham, 2023; Shi et al., 2021). TRV has a bipartite genome (TRV1 and TRV2) with TRV1 containing an annotated movement protein (MP) (**Figure 1A, C**). In previous studies, removal or mutagenesis of the TRV1 MP has rendered the virus incapable of systemic infection (Deng et al., 2013). Conversely, FoMV has a monopartite genome and does not contain an annotated MP but is thought to rely primarily on its coat protein (CP) for systemic infection (**Figure 1D**). Given this information, we sought to immobilize FoMV and TRV by introducing a premature stop codon in the TRV1 MP (MPΔ, **Figure 1B**) after Leu82 and FoMV CP (CPΔ, **Figure 1E**) after Leu189, respectively. Additional sequence introduced during cloning added five amino acids (GRATS) and four amino acids (EGAP) to the C-terminal end of TRV1 MPΔ and FoMV CPΔ, respectively. The plant virus silencing suppressor, P19 was also included on the T-DNA backbone in both immobilized viruses to improve viral replication (**Figure 1B, E**). Furthermore, the fluorescent marker, AmCyan was cloned into multiple cloning sites (MCSs) for each virus to test immobilization, track viral replication and systemic infection (**Figure 1C, E**).

**Figure 1.**
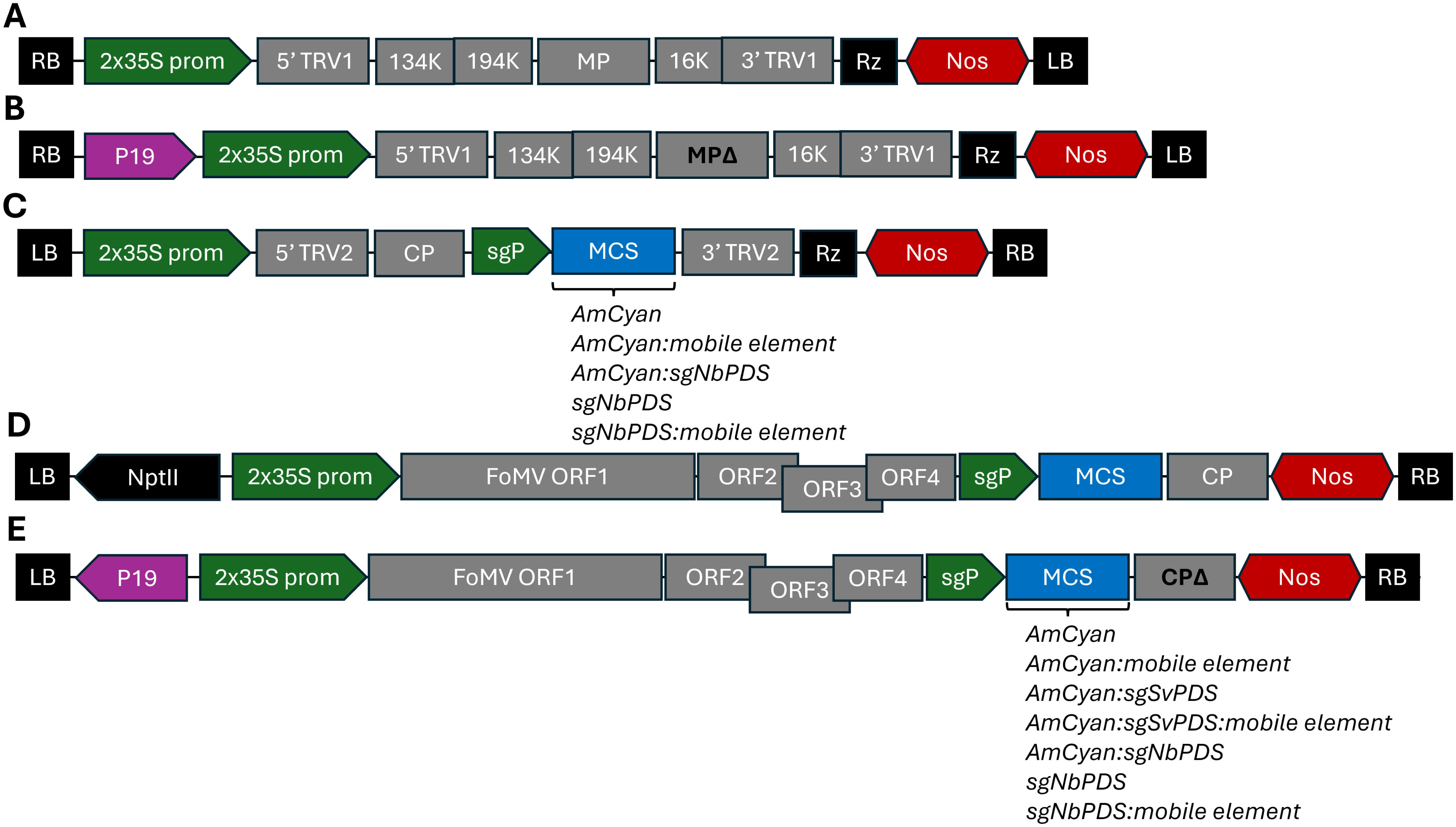
*Tobacco Rattle Virus* (TRV) and *Foxtail Mosaic Virus* (FoMV) T-DNAs used for testing mobile elements for mobility and editing in dicot and monocot species. (**A**) Wild-type TRV1 and (**B**) movement protein (MP) knock-out (MPΔ), immobilized TRV1. (**C**) TRV2 was used for expressing AmCyan and single guide RNA (sgRNA) sequences with and without mobile elements by incorporation into a multiple cloning site (MCS-blue boxes). (**D**) Wild-type FoMV and (**E**) coat protein (CP) knock-out (CPΔ), immobilized FoMV expressing AmCyan and sgRNA sequences with and without mobile elements. TRV1 MPΔ and FoMV CPΔ T-DNAs were generated with and without P19 expression cassettes (purple arrows), with the TRV1 MPΔ P19 cassette incorporating the Nos promoter and EtEU terminator, and the FoMV CPΔ P19 cassette incorporating a 3x enhancer, 35S promoter and 35S terminator. Green arrows indicate promoter or *Pea Early-Browning Virus* sub-genomic promoter (sgP) elements. Gray boxes indicate consecutive and overlapping viral sequences and open reading frames (ORFs). Red hexagons indicate terminator elements. Black boxes indicate right (RB) and left (LB) T-DNA borders. Black arrows indicate selection marker cassettes driven by the 2×35S promoter and 35S terminator. Sequences were incorporated into MCSs using Gibson assembly and AarI and SapI restriction enzyme sites for TRV2 and FoMV, respectively.

### Testing and optimization of immobilized FoMV and TRV viruses using a modified *Nicotiana benthamiana* leaf assay

Agro-infiltration of *Nicotiana benthamiana* (*N. benthamiana*) remains one of the most efficient methods for transient T-DNA delivery and gene expression. In addition to Agrobacterium, *N. benthamiana* is also highly susceptible to a range of both monocot and dicot viruses and can be used to generate so called “viral sap” by processing infected tissues and applying to host plants for infection (Mei et al., 2019). A simple leaf infection assay using *N. benthamiana* leaves was developed using seedlings germinated on media containing the GA-inhibitor, ancymidol (**Figure 2**). Plants germinated on ancymidol are more compact, allowing a complete leaf and leaf vasculature system to be used as an explant for infection and data collection (**Figure 2A**) (Hofmannová et al., 2008). The assay was carried out by cutting away outside edges of each intact leaf, creating a leaf square explant that would present wounded edges for direct Agro-infection and source tissue for viral infection (**Figure 2B**). Assay development was conducted using both wild-type and immobilized FoMV and TRV viruses carrying AmCyan and image data was collected three, five, and seven days post infection (dpi) (**Figures 2** and **S1**).

**Figure 2.**
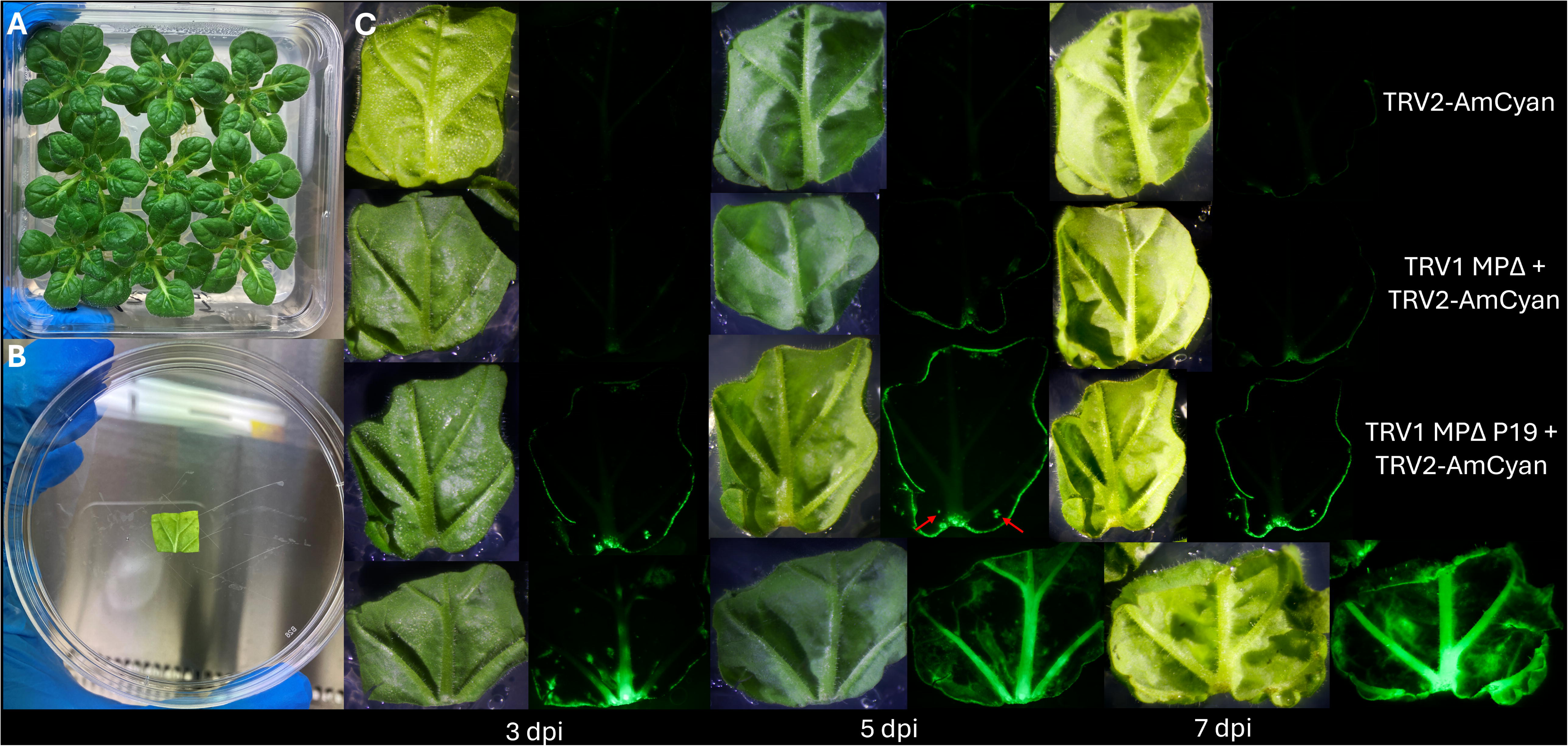
Immobile TRV RNA mobility testing in *Nicotiana benthamiana* (*N. benthamiana*) leaves using AmCyan detection. (**A**) Five- to six-week-old *N. benthamiana* seedlings grown on ancimidol for generating leaf source material. (**B**) Prepared leaf explants used for transient Agrobacterium infection and delivery of TRV T-DNAs (Figure 1). (**C**) Light (first, third, and fifth column of images) and fluorescent (second, forth, and sixth column of images) imaging of leaf explants infected with TRV2-AmCyan only (top row of images; negative control), immobilized TRV1 MPΔ and TRV2-AmCyan (second row of images), TRV1 MPΔ P19 and TRV2-AmCyan (third row of images), and wild-type TRV1 and TRV2-AmCyan (bottom row of images; positive control) T-DNAs three days post infection (3 dpi; first two columns of images), 5 dpi (third and fourth columns of images), or 7 dpi (last two columns of images). Red arrows indicate punctate signal in internal mesophyll tissue from direct wounding.

Overall, TRV treatments showed the strongest viral infection, with wild-type TRV displaying strong signal throughout source, vasculature and mesophyll tissues after 5 dpi (**Figure 2C**) and wild-type FoMV treatments after 7 dpi in all infected explants (**Figure S1**). Immobilized viral treatments with P19 showed stronger infection of source tissues compared to immobile viral treatments without, with complete infection observed in all infected explants for both viruses. Viral infection of immobilized viruses continued for up to two weeks before diminishing while wild-type viruses persisted beyond two-weeks (data not shown). Visual signal of AmCyan in immobile virus source tissues also peaked at 5 dpi and 7 dpi for TRV and FoMV, respectively and was used for subsequent data collection and sampling. P19 expression was necessary for both immobilized viruses to achieve complete infection of source tissues and was used hereafter. Although punctate signal was detected in internal mesophyll of immobilized virus treatments, the majority of signal was detected in source tissues. This background signal is most likely due to direct wounding and Agro-infection of mesophyll tissues from handling explants and appeared stochastically across treatments and controls (**Figure 2C** and **S1**; **red arrows**). Collectively, these data suggest that both immobilized viruses were not capable of systemic infection but in the presence of P19, preserved the ability to replicate and transiently express encoded transcripts effectively in source tissues.

### Visualization and quantification of mobile transcripts using immobilized FoMV and TRV viruses

Previous studies have used immobilized viruses, such as PVX to transiently express mobile transcripts for detection in distal tissues as an alternative to grafting transgenics (Yu et al., 2020). However, these studies were conducted on whole plants and typically do not report direct visualization of mobile transcripts when fluorescent markers were used on this scale. Nevertheless, direct visualization of mobile transcripts has been demonstrated and has been useful for tracking both subcellular and intercellular RNA mobility (Luo et al., 2018). Given this information, we inquired if transient transcript expression from immobilized FoMV and TRV in source tissues would be sufficient for visual detection of mobile transcripts in internal mesophyll tissues in our *N. benthamiana* leaf assay. To test, seven known mobility elements, T-RNA-like sequences (TLSmet, TLSmetΔDT, TLSgly, TLSIle), truncated forms of the Arabidopsis FT transcript (AtFTtruc), and its maize orthologs (ZCN16 and 19) were individually fused to the 3’ end of AmCyan, cloned into the MCS of both immobilized viruses, and tested using the *N. benthamiana* leaf assay (**Figure 3** and **S2**).

**Figure 3.**
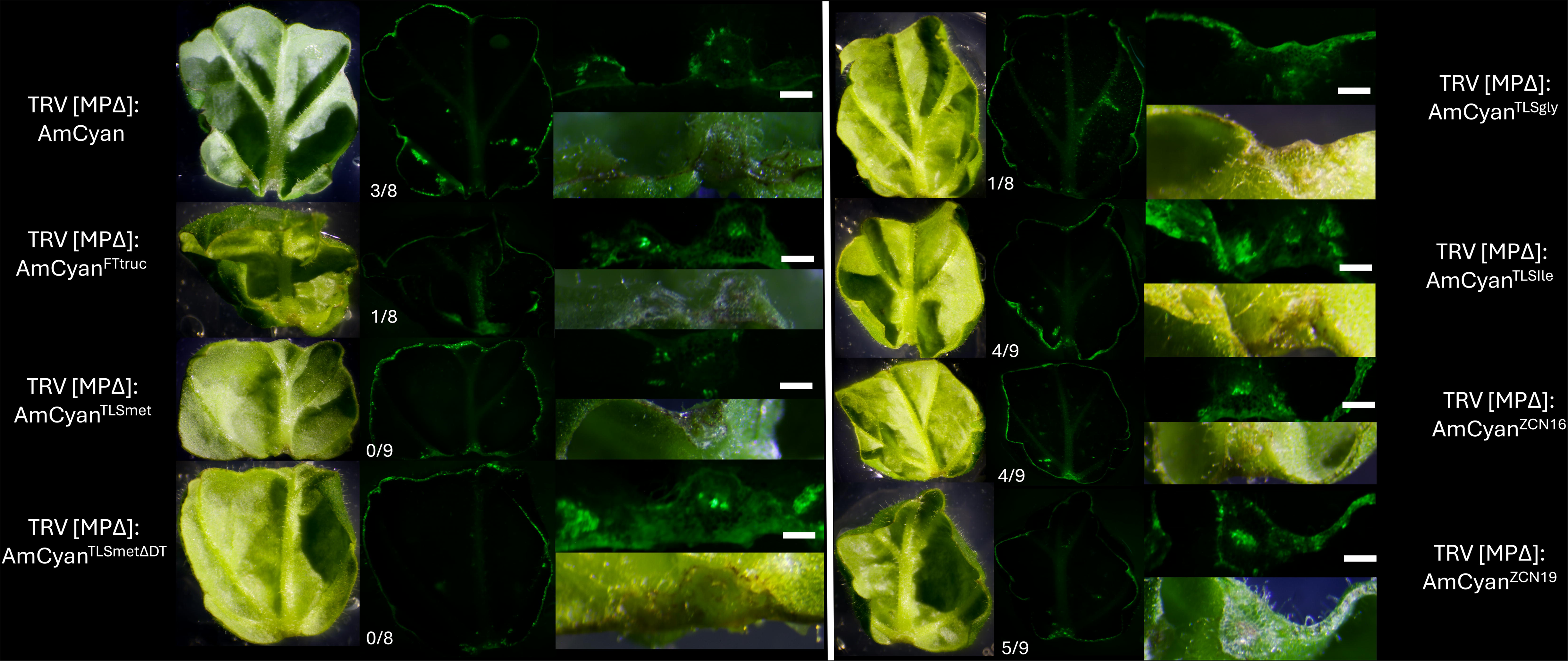
Mobile element testing using immobile TRV and AmCyan detection in *N. benthaminana* leaves. Light (first and fourth column of images) and fluorescent (second and fifth column of images) imaging of leaf explants co-infected with TRV1 MPΔ P19 and TRV2-AmCyan (top left row of images; negative control) or TRV2-AmCyan T-DNAs with AmCyan fused to FTtruc (second left row of images), TLSmet (third left row of images), TLSmetΔDT (bottom left row of images), TLSgly (top right row of images), TLSIle (second right row of images), ZCN16 (third right row of images), and ZCN19 (bottom right row of images) mobile elements. Fractions provide the number of leaf explants with internal AmCyan signal divided by the total number of infected explants. Magnified images of leaf vein cross-sections (third and sixth column of images) with fluorescent (top) and light (bottom) images. Scale bars = 2.5 mm.

Overall, incorporation of the additional mobile element sequence did not seem to destabilize the immobile viruses, with similar infection rates observed in source tissues compared to previous experiments without mobile element sequences (**Figures 2** and **S1**). Furthermore, similar trends with overall signal intensity were observed across both immobile viruses compared to previous experiments without mobile sequences, with overall higher signal detected in source tissues of TRV compared to FoMV. However, internal signals in mesophyll tissues not observed in previous experiments omitting mobile elements were detected consistently within mobile element treatments with varying frequencies across treatments, with some sharing similar frequencies in both viral systems (**Figure 3** and **S2**). Furthermore, source signal detection was strongest in vasculature tissues in mobile element treatments compared to controls without mobile elements, consistent with vascular localization and mobilization (**Figure 2**; **magnified images**). Overall, internal signal detection across treatments was higher for immobile FoMV compared to immobile TRV (**Figure 3** and **S2**). To validate these observations, a hole punch was used to sample the interior of individual leaf explants for RNA extraction and reverse-transcriptase PCR (RT-PCR) of AmCyan and Actin transcripts, Actin acting as a reference transcript (**Figures 4**, **S3** and **S4**).

**Figure 4.**
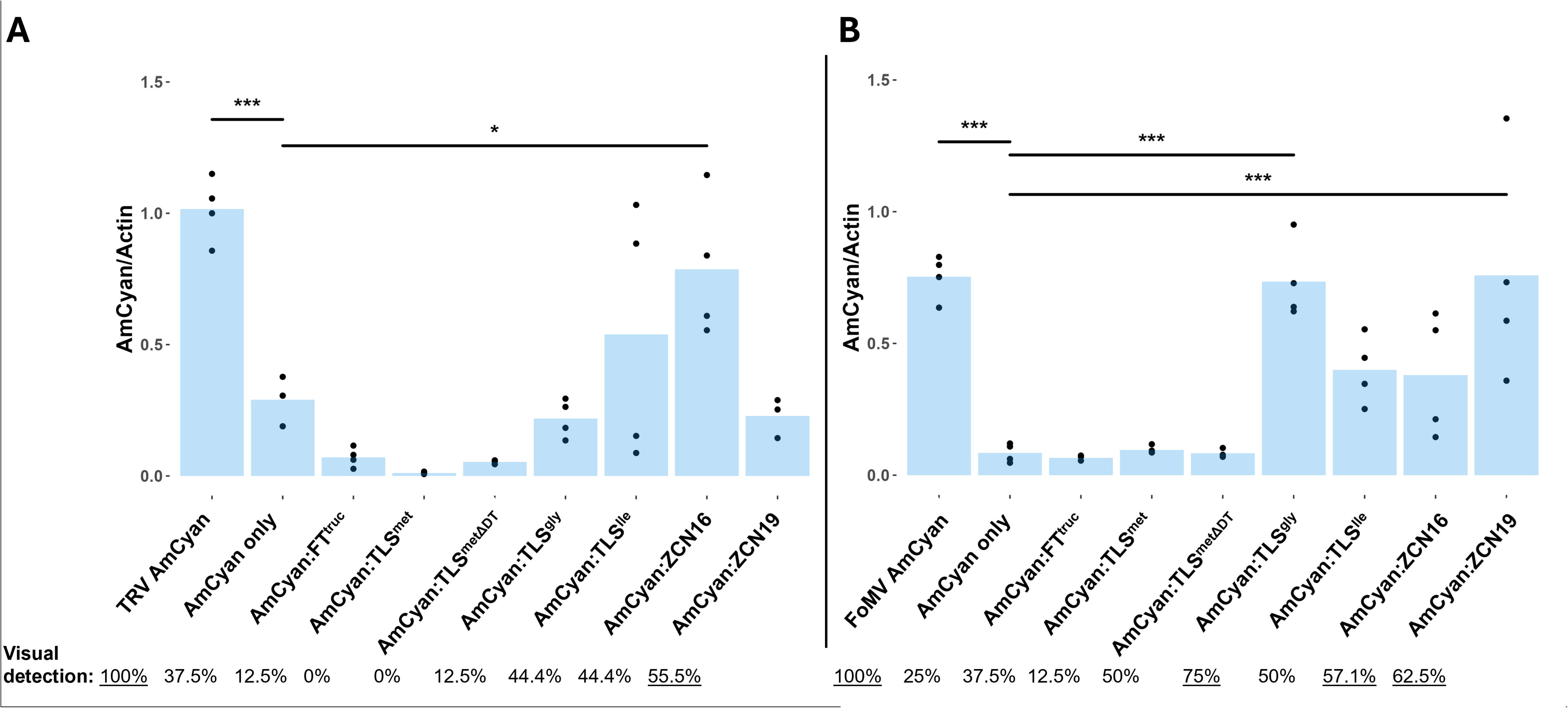
Reverse-transcriptase PCR (RT-PCR) quantification of AmCyan in immobile virus RNA mobility assays. Internal tissues of *N. benthamina* leaves infected with immobilized TRV (**A**; Figure 3) and FoMV (**B**; **Figure S2**) viruses were used for total RNA extraction and RT-PCR of AmCyan and Actin as a reference gene. Gel-based quantification was conducted using Image J (**Figure S3-S4**), calculated as a fraction of AmCyan divided by Actin, and compared to wild-type virus treatments (“TRV AmCyan” and “FoMV AmCyan”). Visual detection percentages (bottom) were calculated using fractions of visual detection of AmCyan signal within internal tissues of leaf explants provided in Figures 3 (**A**) and **S2** (**B**) and underlined if over 50%. Bars are averages of three to four biological replicates and were used to calculate standard error. Black dots are individual data points. Box plot data visualization was used to identify and remove outliers from the analysis falling above 1.5 × IQR from Q3 (no data points fell below 1.5 × IQR from Q1) (**Figures S3-S4**). A Dunnett’s test was used to test significant differences between AmCyan only negative controls and other treatments. *** P≤ 0.001, *P≤ 0.05

Comparing visual signal and RT-PCR data revealed clear trends across mobile element treatments and immobile viruses (**Figure 4**). Control treatments for TRV provided a range from 0.29±0.09 (RT-PCR) and 37.5% (visual) for the AmCyan only negative control to 1.02±0.12 (RT-PCR) and 100% (visual) for wild-type TRV positive control, and a range from 0.08±0.04 (RT-PCR) and 25% (visual) for the AmCyan only negative control to 0.75±0.08 (RT-PCR) and 100% (visual) for the wild-type FoMV positive control. TLSmet and TLSmetΔDT demonstrated the lowest mobility across both immobile viruses with 0.01±0.004 (RT-PCR) and 0% (visual) for TLSmet and 0.05±0.01 (RT-PCR) and 0% (visual) for TLSmetΔDT using immobile TRV, and 0.10±0.01 (RT-PCR) and 12.5% (visual) for TLSmet and 0.08±0.02 (RT-PCR) and 50% (visual) for TLSmetΔDT using immobile FoMV. Unexpectedly, FTtruc also demonstrated low mobility with 0.07±0.04 (RT-PCR) and 12.5% (visual) for immobile TRV, and 0.07±0.01 (RT-PCR) and 37.5% (visual) using immobile FoMV.

Two elements, TLSIle and ZCN16 demonstrated consistent mobility across both viruses, but did not correlate as well between visual and RT-PCR data compared to controls and elements with low mobility (**Figure 4**). For immobile TRV, TLSIle (0.54±0.49, RT-PCR; 44.4% visual) and ZCN16 (0.79±0.27, RT-PCR; 44.4%, visual) demonstrated the highest mobility, while TLSIle (0.40±0.13, RT-PCR; 50%, visual), TLSgly (0.74±0.15, RT-PCR; 75% visual), ZCN16 (0.38±0.24, RT-PCR; 57.1%, visual), and ZCN19 (0.76±0.43, RT-PCR; 62.5%, visual) demonstrated the highest mobility for immobile FoMV. Interestingly, the two elements differing the most between the viruses was TLSgly and ZCN19, which had low mobility for immobile TRV [TLSgly (0.22±0.07, RT-PCR; 12.5%, visual), ZCN19 (0.23±0.08, RT-PCR; 55.6%, visual)] but among the highest mobility for immobile FoMV [TLSgly (0.74±0.15, RT-PCR; 75%, visual), ZCN19 (0.76±0.43, RT-PCR; 62.5%, visual)]. Nevertheless, these data support the use of signal visualization and transcript detection as a proxy for mobile transcript accumulation and suggests TLSIle, TLSgly, ZCN16 and ZCN19 could facilitate mobility of other RNAs.

### Testing optimized mobile elements for virus-induced gene editing in *N. benthimiana* and setaria

Previous studies have fused mobile elements to genome editing reagents to promote RNA stability, mobility and improve editing frequencies with mixed results (Beernink et al., 2022; Nagalakshmi et al., 2022). To test these findings and validate data generated using AmCyan, all seven mobile elements were fused to the 3’ end of a sgRNA targeting the *N. benthimiana PHYTOENE DESATURASE* gene (sgNbPDS) and tested in the *N. benthimiana* leaf assay (**Figure S5**).

Immobile TRV and FoMV viruses were used to test mobile element treatments along with wild-type virus sgRNA only (positive) and immobile virus sgRNA only (negative) controls. Instead of AmCyan signal, phytobleaching resulting from NbPDS knock-out was observed in treated explants, and internal tissue was sampled for genomic DNA and detection of PDS editing (**Figure S5 A-D**). Internal tissues were also sampled for RNA and RT-PCR detection of sgRNAs and Actin as a reference transcript (**Figures S5 E-F**, **S6,** and **S7)**. Despite a reduction in explant replication to support both DNA and RNA analyses, sgRNA detection in TRV showed trends acrosss mobile element treatments relative to the sgRNA only control. Unexpectedly, other than the wild-type positive control, TLSmetΔDT was the only treatment detected above sgRNA only levels for TRV but showed greater variation for FoMV along with other treatments and controls (**Figures S5 E-F**). Furthermore, evidence of photobleaching and edit detection was only observed in wild-type virus positive controls, and immobile FoMV sgRNA only negative control as well as TLSgly and ZCN19 treatments (**Figure S5 A-C**). Although photobleaching and edit detection did not seem to correlate with sgRNA detection, the two mobile elements demonstrating the highest mobility in FoMV AmCyan experiments also produced photobleaching and edits in sgRNA mobility experiments and were used in subsequent FoMV setaria editing experiments (**Figure 5**).

**Figure 5.**
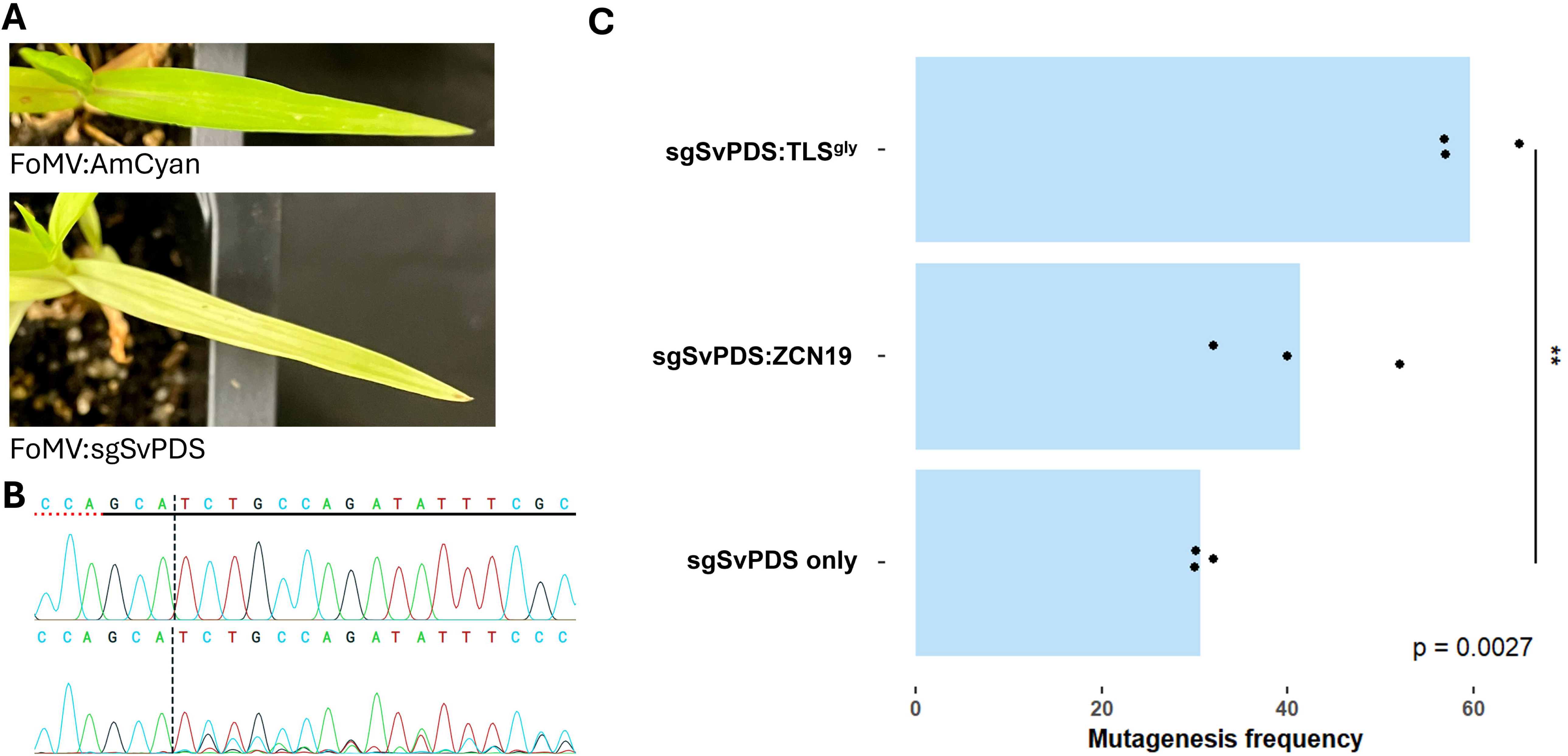
FoMV-mediated targeted mutagenesis in setaria using mobile elements. Setaria seedlings were infected with FoMV carrying AmCyan (FoMV:AmCyan, negative control) or AmCyan fused to sgRNAs targeting the setaria PDS gene (sgSvPDS) with and without mobile elements and systemic leaves were observed for a pale leaf phenotype using light imaging (**A**). Systemic leaves were sampled for total DNA extraction, PCR amplification of the *SvPDS* target site, Sanger sequencing and ICE analysis (Synthego; **B**) to determine mutagenesis frequencies (percentages; **C**). Bars are averages of three biological replicates. Black dots are individual data points. A Dunnett’s test was used to test significant differences between sgRNA only negative controls and other treatments. ** P= 0.0027

As FoMV’s primary host, setaria is highly susceptible to FoMV infection and has been used in previous studies for virus-induced gene silencing (VIGS) (N. Liu et al., 2016; Mei et al., 2016). However, there are limited reports of virus-induced gene editing (VIGE) in setaria (Mei et al., 2019). To test FoMV for VIGE and mobile element function, TLSgly and ZCN19 were fused to the 3’ end of a sgRNA targeting the *Setaria viridis PHYTOENE DESATURASE* gene (sgSvPDS) and tested using wild-type FoMV in setaria seedlings (**Figure 5**). Setaria is highly recalcitrant to Agrobacterium infection. For this reason, viral sap was generated using *N. benthimiana* leaf explants and used for rub-inoculation following previously described methods (Mei et al., 2019). AmCyan was incorporated into all tested viruses to verify viral replication in leaf explants and track viral infection in infected seedlings (**Figure 1**). Wild-type FoMV was used since immobile FoMV was unable to infect setaria seedlings (data not shown). Seedlings were infected with AmCyan only (negative) and sgSvPDS only (positive) controls in addition to mobile element treatments. Seedlings displaying complete systemic infection after two weeks were used to harvest leaf tissue and for edit detection (**Figure 5B**).

A range of editing was observed with controls, ranging from 0% (AmCyan-only, negative) to 30.1%±1.1 (sgRNA only, positive). Both mobile element treatments displayed higher levels of edits than the sgRNA only positive control, with TLSgly displaying a two-fold increase (59.7%±4.6) and ZCN19 displaying a slight increase (41.3%±10.0) (**Figure 5C**). In addition to detected edits, photobleaching was observed in mature somatic tissues of all infected positive control and treatment seedlings (**Figure 5A**). However, no photobleaching was detected in AmyCyan only controls (**Figure 5A**) or germline seedlings sowed from infected seedlings, regardless of viral reagent.

## Discussion

RNA-based technologies are rapidly reshaping biotechnology by enabling programmable control over gene expression, protein translation, and genome editing. Recent advances have focused largely on improving RNA stability, enzymatic activity, and intracellular localization (Jones et al., 2024), while applications in plants have remained dominated by RNA interference and virus-induced gene silencing (VIGS) approaches (Lastovka et al., 2025). The emergence of RNA-guided genome editing and DNA-free transformation strategies has created a growing need for technologies that can systematically evaluate RNA dynamics within plant tissues, particularly long-distance RNA movement, which is increasingly recognized as a key determinant of editing efficiency and heritability (Sharma et al., 2025).

The primary contribution of this study is the development of a simple, generalizable screening platform for evaluating RNA mobility in plants. By immobilizing FoMV and TRV through disruption of proteins affecting viral movement while preserving replication in source tissues, we created viral vectors that uncouple RNA mobility from systemic viral spread. Combined with a simple *Nicotiana benthamiana* leaf assay, these vectors permit rapid visual and molecular assessment of transcript movement without requiring transgenic donor lines, grafting experiments, or intricate vascular sampling. Although grafting has been instrumental for discovering endogenous mobile RNAs, its technical complexity and limited applicability in monocot species have restricted throughput and broader adoption (Thomas & Frank, 2019). Our platform substantially reduces these barriers while providing a generalized experimental framework that can be adapted to diverse RNA cargos and viral systems.

The utility of this platform is demonstrated through screening of previously described RNA mobility elements originating from both dicot and monocot species. Despite differences in viral biology, mobility trends were remarkably consistent between immobilized FoMV and TRV, identifying TLSIle and ZCN16 as reproducibly mobile elements. The differential performance of TLSgly and ZCN19, particularly displaying enhanced activity in FoMV relative to TRV, illustrates that mobility can be influenced by the viral context and host interactions rather than solely by intrinsic RNA sequence. Likewise, the limited mobility observed for TLSmet, TLSmetΔDT and AtFTtruc suggests that previously reported mobile elements may exhibit context-dependent behavior depending on transcript architecture, expression strategy, or experimental system. These observations highlight an important advantage of the platform of being able to compare candidate elements directly under standardized conditions prior to application.

Importantly, screening outcomes translated into functional improvements in viral-mediated genome editing. Mobile elements identified through reporter assays enhanced FoMV-mediated editing in *S. viridis*, with TLSgly producing approximately a two-fold increase in somatic editing efficiency relative to sgRNA alone. While germline editing was not observed under the conditions tested, these results demonstrate that the platform is predictive of biologically meaningful improvements in RNA-mediated editing and can be used to prioritize candidate mobility elements before undertaking more labor-intensive whole-plant experiments. Beyond sgRNAs, the same framework should be readily adaptable for evaluating mobile mRNAs encoding genome editors, recombinases, transcription factors, morphogenic regulators, or other functional RNAs relevant to RNA-based biotechnology.

Several opportunities remain to further expand the platform. Immobilization was achieved by disrupting essential viral movement proteins, which also prevented systemic infection of whole plants. Future iterations could employ more subtle amino acid substitutions, conditional mutations, or inducible movement functions to preserve broader infectivity while maintaining spatial control over viral spread (Dawson & White, 1979; Garcia-Ruiz, 2018). Likewise, extending this strategy to additional viral families would broaden the range of hosts and applications available for comparative studies of RNA mobility.

Monocot crops, including sorghum (*Sorghum bicolor*) and maize (*Zea mays*), remain among the most important agricultural species world-wide but continue to lag behind dicots in the availability of RNA-based engineering tools (Singh et al., 2025). Efficient evaluation of mobile RNAs has been particularly challenging in these species because grafting is difficult and viral delivery systems remain limited (Steinberger & Voytas, 2025). By providing a rapid, modular, and experimentally accessible assay for screening RNA mobility elements, immobilized FoMV and TRV vectors establish a foundation for engineering mobile RNAs across diverse plant systems. As RNA-guided genome editing, DNA-free transformation, and mobile RNA technologies continue to develop, this scalable screening platform should accelerate discovery of functional RNA elements and facilitate the development of next-generation RNA delivery systems for both fundamental research and crop improvement.

## Experimental Procedures

### Vector construction and Agrobacterium culture preparation

Viral T-DNA vectors were constructed using Gibson assembly and the FoMV cloning vector, pEE082 (Baysal *et al*., 2024) and the TRV1 and TRV2 cloning vectors, pYL192 (Addgene #148968) and SPDK3876 (Addgene #149275), respectively (Ellison et al., 2020). **Table S1** contains vector backbones, linearizing restriction enzymes, and fragments used for Gibson assembly. Fragments were either synthesized (ThermoFischer Scientific) or generated using Chamness et al., 2023 components as templates for PCR using the Q5 DNA polymerase (New England Biolabs). Gibson assembly was conducted using the Gibson Assembly® Master Mix (New England Biolabs) with recommended conditions and 75 ng linearized vector. Vectors were transformed into GV3101 Agrobacterium and grown on LB media containing kanamycin (50 mg/L) and gentamicin (30 mg/L).

### Plant materials

*Nicotiana benthamiana* (*N. benthamiana*) expressing Cas9 was used for *N. benthimiana* leaf infection assays and provided from a previous study (Ellison et al., 2020). *N. benthamiana* seed was generated by growing plants in growth chambers at 25°C and a photoperiod of 16/8 h day/night. *N. benthamiana* seed was sterilized using 15% bleach for 20 mins, washed three times with sterile deionized water, and germinated on MS in a growth incubator at 25°C with a photoperiod of 16/8 h day/night. MS media contained 2.165 g/L MS basal salts (Sigma M5524), 20 g/L sucrose (PhytoTech S829), 5 g/L plant Agar (Sigma A7921), 3.4 mg/L ancymidol (Sigma A9431), 5 mg/L meropenem trihydrate (PhytoTech M5600) with pH 5.6 using NaOH. Three- to four-week-old seedlings were transferred to Phytatrays^TM^ (Sigma-Aldrich) containing MS media for preparation of leaf source material.

*Setaria viridis* (setaria) expressing Cas9 was used for setaria virus infection experiments and provided from a previous study (Mei et al., 2019). Setaria seed was generated by growing plants in growth chambers at 28°C and a photoperiod of 16/8 h day/night. Setaria seed was sterilized using 15% bleach for 20 mins, washed three times with sterile deionized water, and germinated on MS in a growth incubator at 25°C with a photoperiod of 16/8 h day/night. A twenty-four hour room temp incubation period in 5% liquid smoke was used to break dormancy if needed. Two- to three-week-old seedlings were transferred to soil and grown under diurnal conditions (22°C day/16°C night) with a photoperiod of 16/8 h day/night in growth chambers. Seedlings were infected one or two-weeks post transfer to soil.

### *N. benthimiana* leaf infection assay

Fully opened leaves of non-flowering *N. benthimiana* plants were used as source material for *N. benthimiana* leaf infection assays using Agrobacterium (**Figure 2A**). Four-to-eight leaves were harvested per treatment and four cuts were made with a scalpel blade to remove leaf margins (**Figure 2B**). Prepared leaf explants were transferred to sterile petri dishes containing Infiltration buffer. Infiltration buffer contained 1.95 g/L MES (PhytoTech M825), 2.03 g/L MgCl_2_·6H□O (Sigma M2393), 200 µg acetosyringone (PhytoTech A104) with pH 5.6 using NaOH. Once explant preparation was complete, Infiltration buffer was replaced with an Agrobacterium solution prepared using Agrobacterium cultures diluted to ∼0.5-0.6 OD600 using Infiltration buffer and co-incubated for ∼20 min with gentle agitation. Post co-incubation, excess solution was gently shaken off, explants were transferred to Co-culture media adaxial face down, and co-cultured for 72 hrs at 25°C in the dark. Co-culture media contained 2.165 g/L MS basal salts (Sigma M5524), 30 g/L sucrose (PhytoTech S829), 8 g/L plant Agar (Sigma A7921), 0.8 mg/L BAP (PhytoTech B800), 0.1 mg/L IBA (PhytoTech I538), 200 µg acetosyringone (PhytoTech A104) with pH 5.6 using NaOH. Post co-culture, explants were transferred to Resting media and allowed to rest in growth incubators at 25°C and a photoperiod of 16/8 h day/night prior to data collection, sampling, or viral sap preparation following methods from (Mei et al., 2019). Resting media contained the same components as Co-culture media, omitting acetosyringone and including 50 mg/L meropenem trihydrate (PhytoTech M5600).

### Setaria virus infection experiments

Fully emerged leaves of non-flowering setaria seedlings were infected as source tissue using viral sap generated from *N. benthimiana* leaf infection assays in virus infection experiments. Viral sap was collected one-to-two weeks post co-culture and processed using previously described methods (Mei et al., 2019). Rub inoculation was carried out by rubbing leaf sap containing silicon carbide (Sigma 357391) directly onto source leaves. Inoculated seedlings were grown under diurnal conditions (22°C day/16°C night) with a photoperiod of 16/8 h day/night in growth chambers and observed for AmCyan viral tracking one-to-two weeks post inoculation. Seedlings displaying complete infection were used for data collection and seed harvesting.

### Detection of AmCyan fluorescence and *in planta* viral RNA tracking

*N. benthimiana* leaf explants used in the *N. benthimiana* leaf infection assays were visualized for AmCyan fluorescence to 1) track viral replication in infected source tissues, 2) track systemic viral mobility (wild-type viruses) or score mobile RNA (immobile viruses) in internal tissues, or 3) track viral replication in infected leaf explants for viral sap preparation. Setaria seedlings used in viral infection experiments were visualized for AmCyan fluorescence to track viral infection for data collection and sampling. AmCyan fluorescence was detected using the Xite Fluorescence Flashlight System (Electron Microscopy Sciences, PA, USA) and methods previously described (Butler et al., 2025).

*N. benthimiana* leaf explants were selected for data collection, sampling and viral sap preparation only if complete viral infection of source tissues was observed and sampled for DNA and RNA extraction using an 8 mm diameter hole punch made in the center of the explant as individual samples. Setaria seedlings were selected for data collection and sampling only if complete viral infection of systemic tissues were observed. *N. benthimiana* leaf explants were scored positive for internal signal if any signal was observed in internal tissues.

### RT-PCR, PCR and edit detection analysis

Total RNA for RT-PCR analysis was extracted using the RNeasy Plant Mini Kit following recommended conditions (Qiagen # 74904). The OneStep RT-PCR kit (Qiagen # 210212) was used for all RT-PCR experiments under recommended conditions for 25 µL reactions and oligos listed in **Table S2**. For AmCyan detection, 50 ng total RNA, 65°C annealing temp, 1 min extension time and 25 cycles were used. For sgRNA detection, 250 ng total RNA, 61°C annealing temp, 15 s extension time and 40 cycles were used. For actin detection, 50 ng total RNA, 62°C annealing temp, 15 s extension time and 35 cycles were used and primers from Liu et al., 2012. Complete RT-PCR reactions were run on 2.0% agarose gels and bands were quantified using ImageJ.

Total genomic DNA for PCR analysis and edit detection was extracted using the DNeasy Plant Pro Kit following recommended conditions (Qiagen # 69204). 150 ng total genomic DNA was used for PCR using the Q5 DNA polymerase (New England Biolabs # M0491), recommended conditions for 50 µL reactions, and oligos listed in **Table S2**. Cycling conditions for all PCR experiments included a 64°C annealing temp, 30s extension time and 32 cycles were used. Edit detection analysis of PCR products was conducted using Sanger sequencing (Eurofins Genomics LLC) and indel frequency was quantified using the ICR CRISPR Analysis Tool (Conant et al., 2022).

## Author contributions

N.M.B conceived the research, conducted the experiments, and wrote the manuscript. C.M.G. assisted with creating graphs and statistical analysis. C.S. assisted in vector design and cloning.

## Acknowledgements

We thank Dan Voytas and his lab for assistance in providing source materials for TRV and setaria virus infection experiments, and Steve Whitham for providing FoMV viral vectors. Funding for this work was provided by the Department of Energy Biological and Environmental Research (BER) Program (US Department of Energy, Award Number DE SC0023160) and Department of Energy Center for Advanced Bioenergy and Bioproducts Innovation (CABBI) (US Department of Energy, Award Number DE SC0018420).

## Supporting information

**Figure S1.** Immobile FoMV RNA mobility testing in *N. benthamiana* leaves using AmCyan detection.

**Figure S2.** Mobile element testing using immobile FoMV and AmCyan detection in *N. benthaminana* leaves.

**Figure S3.** Reverse-transcriptase PCR (RT-PCR) gel images used for quantification of AmCyan in immobile TRV RNA mobility assays.

**Figure S4.** Reverse-transcriptase PCR (RT-PCR) gel images used for quantification of AmCyan in immobile FoMV RNA mobility assays.

**Figure S5.** Mobile element testing using sgRNA detection and CRISPR mutagenesis via N. benthamina immobile virus assay.

**Figure S6.** Reverse-transcriptase PCR (RT-PCR) gel images used for quantification of sgNbPDS in immobile TRV RNA mobility assays.

**Figure S7.** Reverse-transcriptase PCR (RT-PCR) gel images used for quantification of sgNbPDS in immobile FoMV RNA mobility assays.

**Table S1.** Backbones and fragments for Gibson assembly

**Table S2.** Oligos for PCR and RT-PCR

