## Supplemental Figures for "Immobilized dicot and monocot viral vectors enable rapid screening of RNA mobility elements for mobile RNA engineering and RNA-based genome editing"

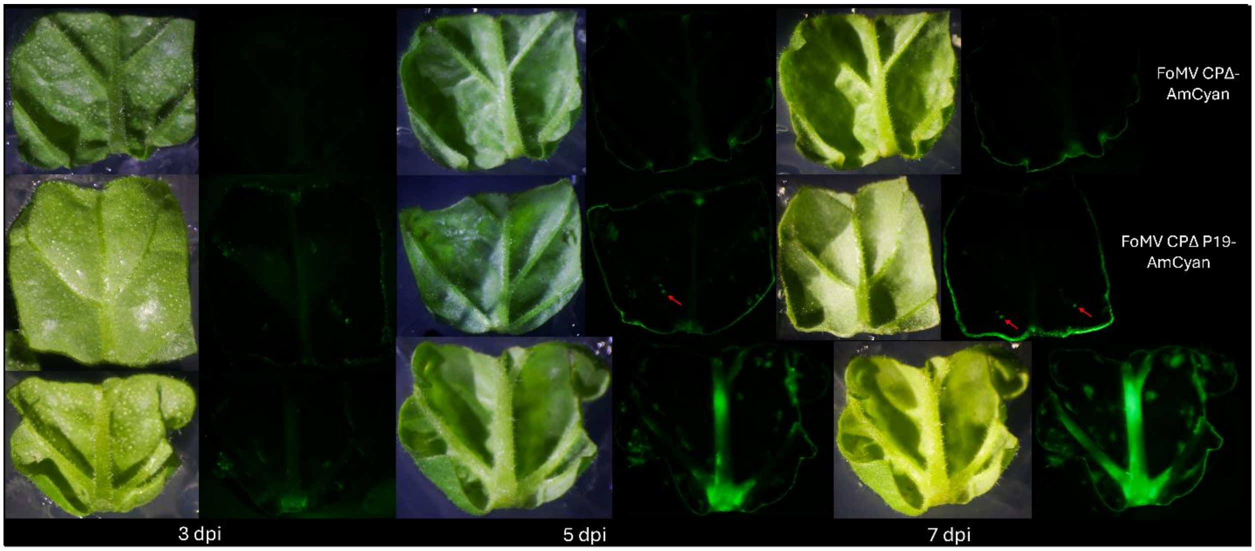


**Figure S1.** Immobile FoMV RNA mobility testing *in Nicotiana benthamiana* (*N. benthamiana*) leaves using AmCyan detection. Light (first, third, and fifth column of images) and fluorescent (second, forth, and sixth column of images) imaging of leaf explants infected with immobilized FoMV CPΔ-AmCyan (top row of images; negative control), FoMV CPΔ P19 (second row of images), and wild-type FoMV-AmCyan (bottom row of images; positive control) T-DNAs three days post infection (3 dpi; first two columns of images), 5 dpi (third and fourth columns of images), or 7 dpi (last two columns of images). Red arrows indicate punctate signal in internal mesophyll tissue from direct wounding.

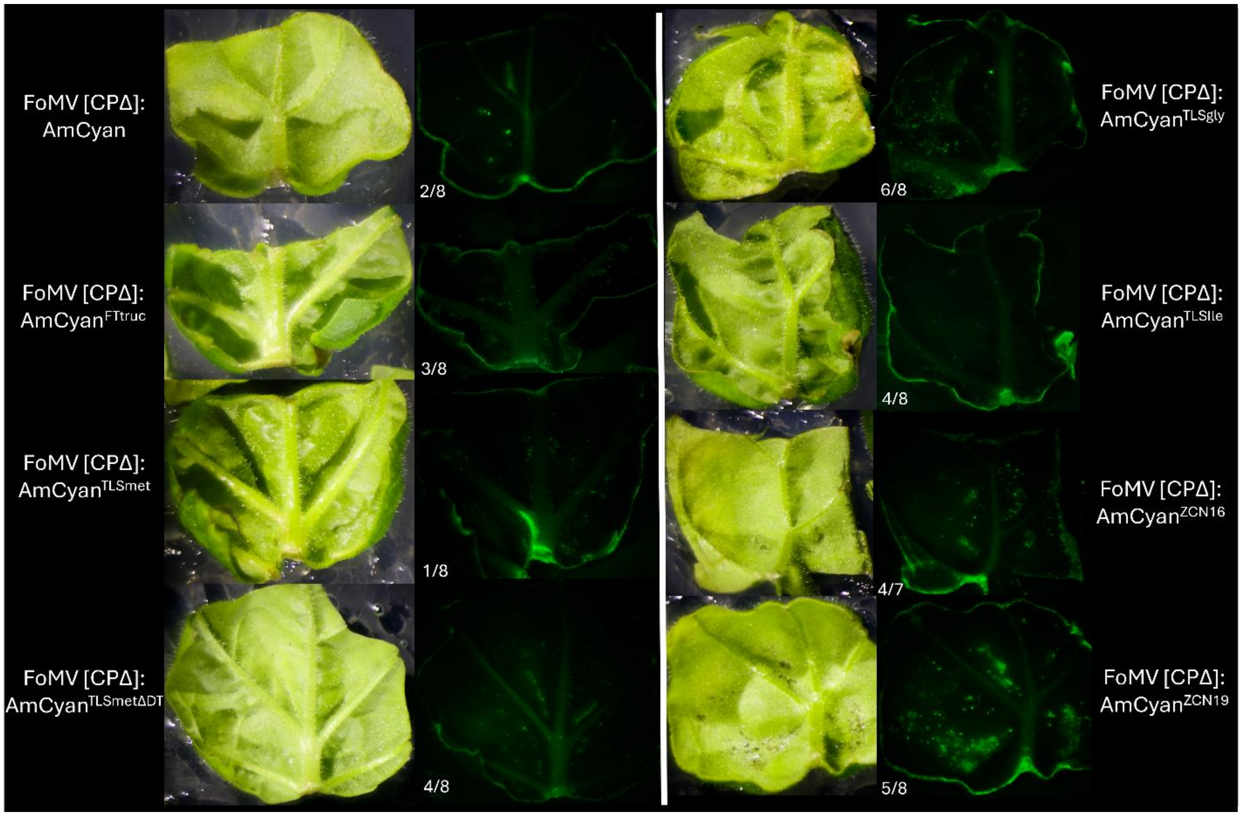


**Figure S2.** Mobile element testing using immobile FoMV and AmCyan detection in *N. benthaminana* leaves. Light (first and third column of images) and fluorescent (second and fourth column of images) imaging of leaf explants infected with FoMV [CPΔ] P19-AmCyan (top left row of images; negative control) or FoMV [CPΔ] P19-AmCyan T-DNAs with AmCyan fused to FTtruc (second left row of images), TLSmet (third left row of images), TLSmetΔDT (bottom left row of images), TLSgly (top right row of images), TLSIle (second right row of images), ZCN16 (third right row of images), and ZCN19 (bottom right row of images) mobile elements. Fractions provide the number of leaf explants with internal AmCyan signal divided by the total number of infected explants.

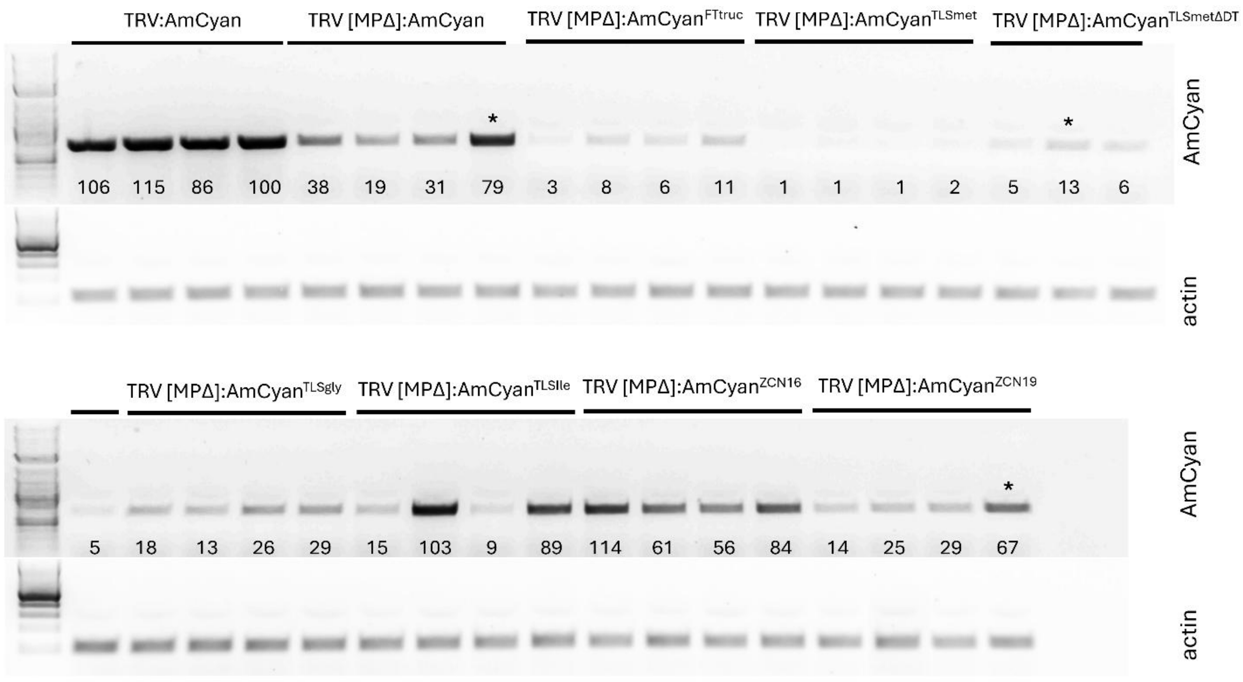


**Figure S3.** Reverse-transcriptase PCR (RT-PCR) gel images used for quantification of AmCyan in immobile TRV RNA mobility assays. Internal tissues of *N. benthamina* leaves infected with immobilized TRV were used for RT-PCR of AmCyan (top gel images) and Actin (bottom gel images) using primers provided in **Table S2**. Gel-based quantification was conducted using Image J (values; AmCyan) for four biological replicates. 2-log DNA ladder was used to determine size of the AmCyan (690 bp) and Actin (114 bp) amplicons. Asterisks indicate identified outliers removed from the analysis (**Figure 4**).

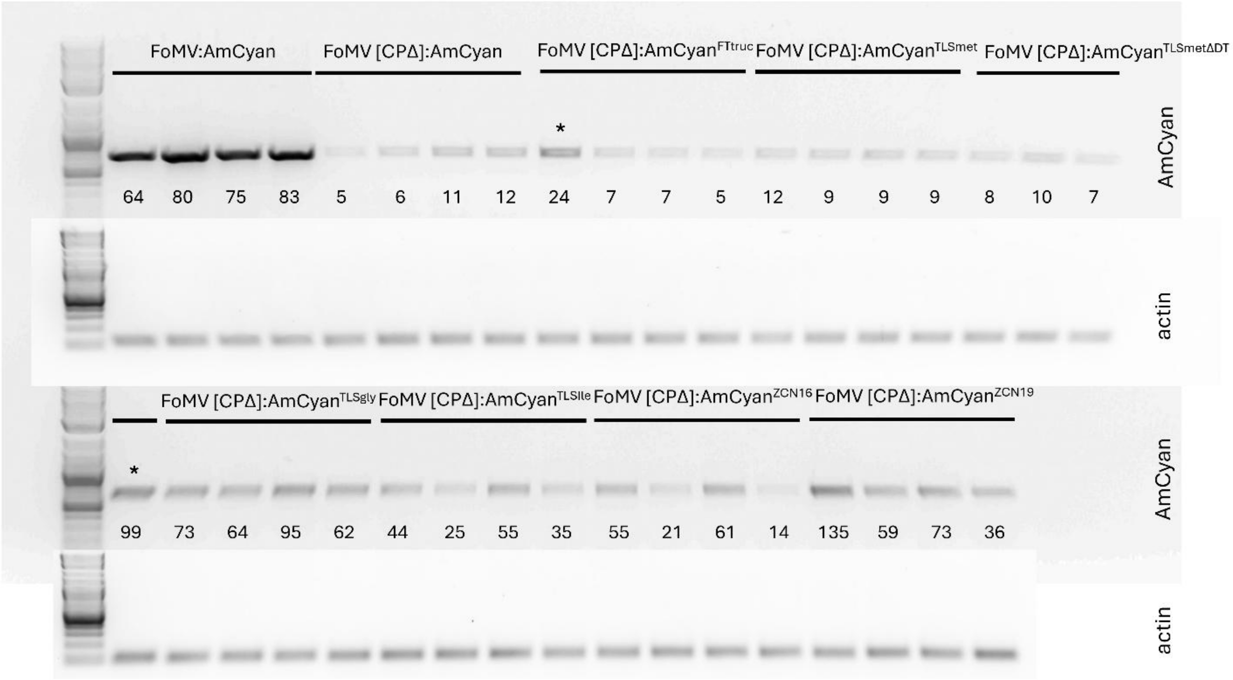


**Figure S4.** Reverse-transcriptase PCR (RT-PCR) gel images used for quantification of AmCyan in immobile FoMV RNA mobility assays. Internal tissues of *N. benthamina* leaves infected with immobilized FoMV were used for RT-PCR of AmCyan (top gel images) and Actin (bottom gel images) using primers provided in **Table S2**. Gel-based quantification was conducted using Image J (values; AmCyan) for four biological replicates. 2-log DNA ladder was used to determine size of the AmCyan (690 bp) and Actin (114 bp) amplicons. Asterisks indicate identified outliers removed from the analysis (**Figure 4**).

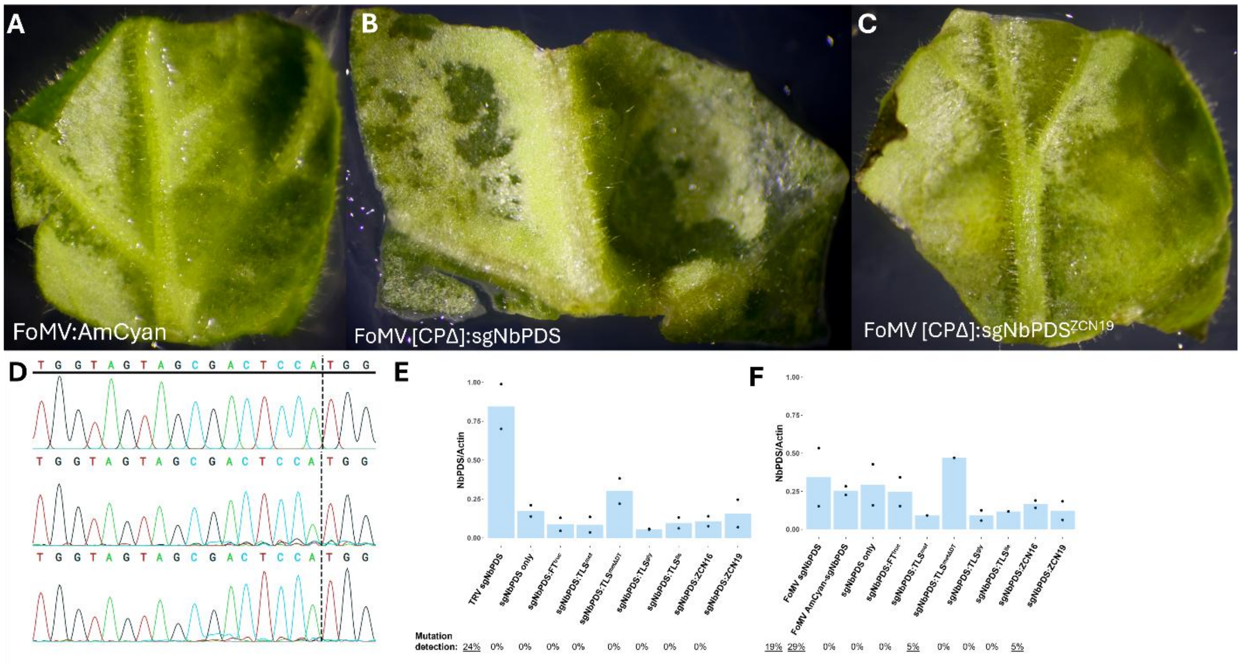


**Figure S5.** Mobile element testing using sgRNA detection and CRISPR mutagenesis via *N. benthamina* immobile virus assay. Light images of leaf explants infected with wild-type FoMV-AmCyan (**A**; negative control) or immobilized FoMV (**B**-**C**) and TRV (not shown) expressing a sgRNA targeting the *N. benthamina* PDS gene (sgNbPDS; **B**) or sgNbPDS fused to a mobile element (**C**; example ZCN19). Internal tissues were used for total DNA extraction, PCR amplification of the *NbPDS* target site using primers provided in **Table S2**, Sanger sequencing, and ICE analysis (Synthego; **D**) to determine mutagenesis frequencies (percentages; **E**). Internal tissues were also used for total RNA extraction and RT-PCR of sgNbPDS and Actin as a reference gene. Gel-based quantification for immobile FoMV (**E**) and TRV (**F**) was conducted using Image J (**Figures S6**-**S7**), calculated as a fraction of sgNbPDS divided by Actin, and compared to wild-type virus treatments ("FoMV AmCyan-NbPDS" and "TRV AmCyan-NbPDS"). Bars are averages of two biological replicates. Black dots are individual data points. Box plot data visualization was used to identify and remove outliers from the analysis falling above 1.5 × IQR from Q3 (no data points fell below 1.5 × IQR from Q1) (**Figures S6-S7**). A Dunnett's test was used to test significant differences between AmCyan only negative controls and other treatments.

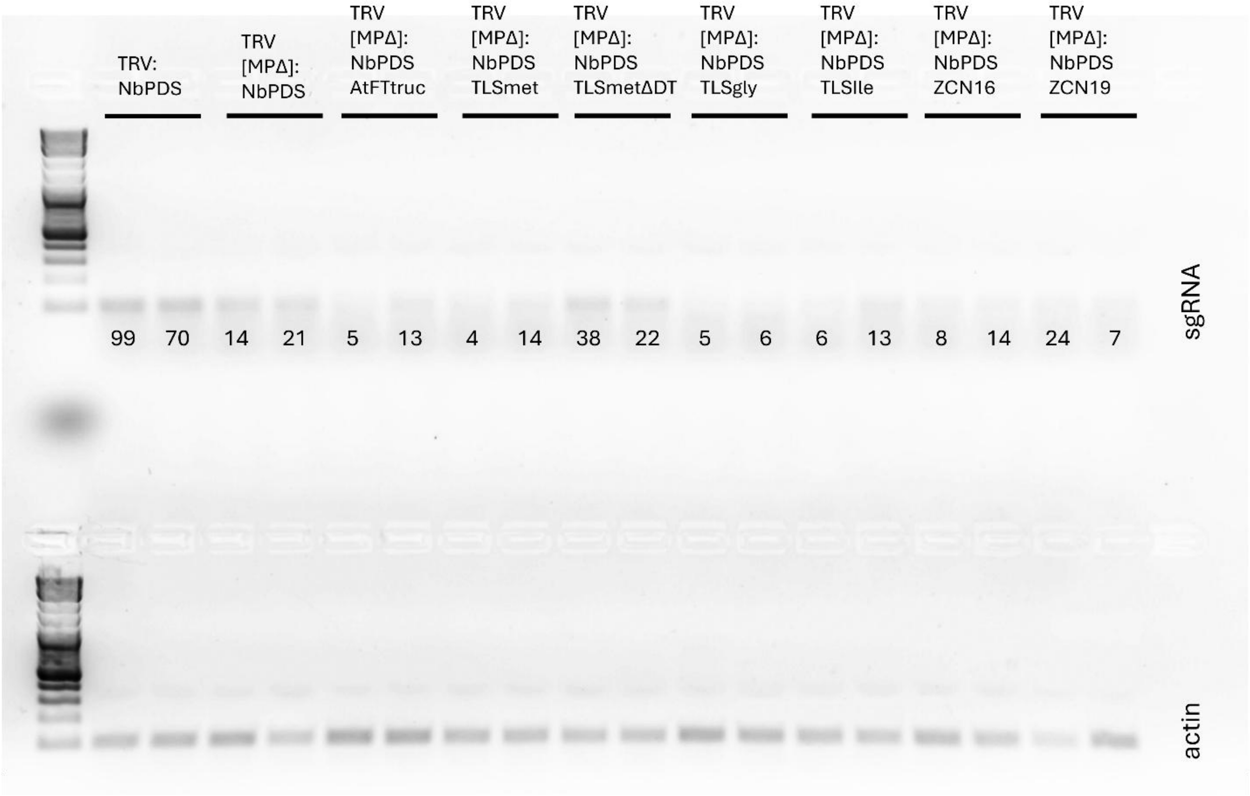


**Figure S6.** Reverse-transcriptase PCR (RT-PCR) gel images used for quantification of sgNbPDS in immobile TRV RNA mobility assays. Internal tissues of *N. benthamina* leaves infected with immobilized TRV were used for RT-PCR of sgNbPDS (top gel images) and Actin (bottom gel images) using primers provided in **Table S2**. Gel-based quantification was conducted using Image J (values; AmCyan) for two biological replicates. 2-log DNA ladder was used to determine size of the sgNbPDS (106 bp) and Actin (114 bp) amplicons.

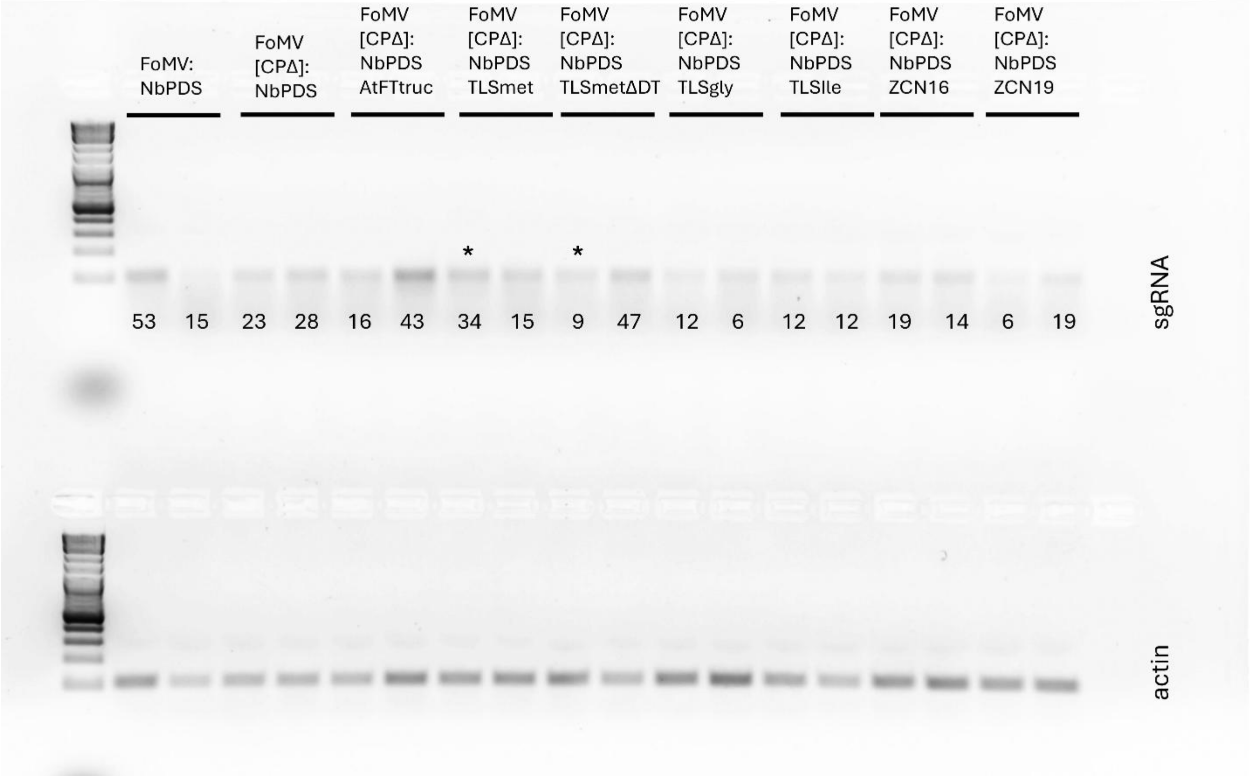


**Figure S7.** Reverse-transcriptase PCR (RT-PCR) gel images used for quantification of sgNbPDS in immobile FoMV RNA mobility assays. Internal tissues of *N. benthamina* leaves infected with immobilized FoMV were used for RT-PCR of sgNbPDS (top gel images) and Actin (bottom gel images) using primers provided in **Table S2**. Gel-based quantification was conducted using Image J (values; AmCyan) for two biological replicates. 2-log DNA ladder was used to determine size of the sgNbPDS (106 bp) and Actin (114 bp) amplicons. Asterisks indicate identified outliers removed from the analysis (**Figure S5**).
